# Brain Stroke Prediction: A PSO Optimized Stacked Ensemble Machine Learning Approach with SMOTEEN Data Balancing

**DOI:** 10.1101/2025.06.05.657994

**Authors:** Achin Jain, Arun Kumar Dubey, Meenakshi Gupta, Sarita Yadav, Arvind Panwar, Roger Atanga, Bernardo Lemos, Saurav Mallik

**Affiliations:** Department of Information Technology, Bharati Vidyapeeth’s College of Engineering, New Delhi; Department Of Electronics And Communications, Manav Rachna University, Sector-43, Aravali Hills, Surajkund Road, Faridabad, Haryana-121001; Galgotias Multi-Disciplinary Research Development Cell(G-MRDC)), Galgotias University, Greater Noida-201308,UP(India); Department of Environmental Health, Harvard T H Chan School of Public Health, Boston, MA, 02115, USA; Department of Pharmacology & Toxicology, University of Arizona, Tucson, AZ, 85721, USA

**Keywords:** Stroke prediction, Machine learning, Stacking ensemble, Class Imbalance, SMOTEEN, PSO Optimization

## Abstract

Prediction of stroke is a critical challenge in healthcare, where early intervention can significantly reduce risks and improve patient outcomes. Traditional methods often struggle with imbalanced datasets and low prediction accuracy. This paper proposes a novel approach to address these issues by combining PSO-optimized Stacked Ensemble ML Classifiers with SMOTEEN data balancing to predict strokes effectively. The technique optimizes 3 Baseline ML Classifiers(RF, SVM, XGB) using Particle Swarm Optimization (PSO) and Stacked Ensemble approach is applied that significantly improving prediction performance. The proposed methodology achieved an impressive 95.57% accuracy, outperforming conventional models such as SVM, LR, RF, KNN, XGB. Overall, the PSO-optimized stacked ensemble approach outperforms traditional machine learning techniques in stroke prediction, providing a more accurate and robust model for early detection and intervention.

## I. Introduction

Stroke is a serious medical condition and a leading cause of death worldwide, occurring when blood flow to the brain is disrupted. According to the World Stroke Organization, one in four people over 25 is at risk of experiencing a stroke. The widespread impact of the condition underscores the need for accurate and timely prediction methods to improve patient outcomes. As the world population has grown, there has been an alarming rise in brain strokes, which will result in a significant rise in annual deaths by 2023. The need to solve this situation has become more urgent as the number of stroke-related deaths continues to increase. Stroke research is now at the forefront of medical research due to this concerning trend.

By examining large data sets that include demographic data, medical history, and physiological markers such as age, blood pressure, and glucose levels, machine learning algorithms have shown promise in transforming the prediction of strokes [1], [2]. However, there are issues that need to be resolved when implementing these algorithms in clinical contexts. Potential bias in training data is an alarming concern, as it can lead to biased predictions and unequal healthcare outcomes [3]. Disparities in healthcare access, socioeconomic considerations, and insufficient or unrepresentative datasets can all lead to biases. Ensemble learning techniques such as stacking have become a reliable way to address these issues. By integrating several classifiers, stacking might enhance the predictive system’s overall generalization capacity and lessen the effects of biases present in individual models.

Furthermore, principal component analysis (PCA) is a powerful dimensionality reduction method. It helps simplify data representations by linearly converting the original features into orthogonal features called principal components. These are arranged according to the variance. This allows PCA to reduce complex datasets into a lower-dimensional space. In this process, it preserves important features.PCA determines the directions of highest variance and their corresponding magnitudes by identifying eigenvectors and eigenvalues from the data covariance matrix [4]-[6]. PCA is used in many fields, such as feature extraction, noise reduction, and data visualization [7]. The suggested research has achieved remarkable predictive accuracy, reaching an astounding 95.57%, through a groundbreaking approach for predictive analysis in ischemic brain stroke using sophisticated machine learning techniques, such as various ML algorithms and ensemble learning procedures.

Because it provides a means of improving prediction accuracy by combining several classifiers, ensemble learning has gained attention in the domains of machine learning and computational intelligence. Ensemble approaches were first developed to increase classification accuracy, but have since been developed to address a variety of real-world problems, including adjusting to shifting concepts, fixing mistakes, choosing the most pertinent characteristics, gradually learning and assessing confidence levels. Significant progress has been made in recent years as a result of the in-depth exploration by researchers of different fusion strategies and the elements that comprise the ensembles [8]-[11]. Stroke prediction is a complex field that requires careful consideration of challenges such as accuracy, missing data, data imbalance, and interoperability. However, A novel approach can utilize the potential of machine learning. The main highlights of proposed worked are here.

### A. Key Contributions

1. To Conduct a background study and provide a review of the literature on brain stroke detection, highlighting the existing techniques used for the assessment of stroke risk.
2. To propose a novel methodology **PSO Optimized Stacked Ensemble ML Classifier using SMOTEEN Data Balancing** to address the problem of data imbalance in stroke prediction. The novelty aspect of this technique lies in using Particle Swarm Optimization (PSO) to optimize 3 Baseline ML Classifiers(RF, XGB, SVM) and then applying the stacking ensemble model for data balancing.
3. To test and validate the proposed methodology using various performance metrics, such as precision, precision (p), recall (r), F1 score, AUC, and AUCPR, ensuring a comprehensive evaluation and robustness of the model.
4. To compare the performance of the proposed methodology against baseline machine learning classifiers, as well as other optimization techniques like Bayesian Optimization (BO), Differential Evolution (DE), and Ant Colony Optimization (ACO), to demonstrate the improvements and benefits of the proposed approach.

The remainder of the paper is organized as follows. Section II reviews the Related Work on stroke prediction models. Section III describes the materials and methods, including the data set, pre-processing, data balancing using SMOTEEN, and optimization with PSO. In Section IV, we present the Proposed System Architecture. Section V outlines the Experimental Setup and evaluation metrics. Section VI discusses the results and discussions, comparing the proposed model with existing methods. Finally, Section VII concludes the paper and suggests future work.

## II. Related Work

The application of machine learning (ML) techniques has significantly advanced stroke prediction research, leading to improved accuracy and timely interventions. This literature review synthesizes recent studies on ML-based stroke prediction, focusing on algorithmic performance, feature selection methods, model interpretability, and key outcomes. In [12], a novel stroke detection algorithm integrates classifiers such as Naïve Bayes, logistic regression, XGBoost, and support vector machines (SVM). The SVM algorithm demonstrated superior performance, achieving 98.6% accuracy and 99.9% precision.

However, the study lacked detailed discussions on feature selection and data pre-processing techniques. [13] explores an algorithm based on ML that uses hospital presentation data and social determinants of health (SDoH). Inclusion of SDoH variables markedly improved sensitivity, specificity, and AUC scores, increasing from 0.694 to 0.823. This study highlights the consistent outperformance of ML classifiers over logistic regression.

The authors in [14] utilized various datasets and machine learning models, including Naive Bayes, SVM, Decision Tree, Random Forest, and Logistic Regression. This study introduces the minimal genetic folding model (MGF), which achieves 83. 2% precision and outperforms other models in AUC scores. Although the MGF model shows promise in distinguishing stroke recovery stages, more research is needed. A potential limitation is the oversampling method, which may have impacted the classifier’s performance.

In [15], logistic regression with recursive feature elimination (RFE) was applied to predict stroke and transient ischemic attack (TIA), focusing on the symptoms reported by the patient. The models achieved an AUC greater than 0.94 and performance was further enhanced by incorporating follow-up data. [16] highlights the efficacy of stack classification, combining base classifiers such as Nave Bayes and random forests with a logistic regression metaclassifier. This method achieved an AUC of 98.9% and 98% accuracy, demonstrating its robustness across multiple performance metrics. [17] examines the interpretability of the model for stroke prediction using SHAP and LIME. Random Forest emerged as the model with the best performance with 90. 36% accuracy, closely followed by the XGB classifier at 89.02%. [18] applies ML to predict ischemic stroke in emergency settings, focusing on clinical laboratory features. The XGBoost-based model stood out, demonstrating the highest accuracy for ischemic stroke prediction and consistent validation across multiple datasets. The high sensitivity and specificity of the model further affirm its reliability in prescreening patients.

The authors in [20] propose a stroke prediction strategy using logistic regression, achieving 86% precision through techniques such as SMOTE, feature selection, and outlier control. Factors such as blood pressure, age, glucose levels, and previous stroke history are analyzed. Although logistic regression outperforms other models, Naive Bayes achieves 82% accuracy in related studies. The research highlights the potential of machine learning for early stroke detection and prevention. In [21] the proposed model achieves 94% accuracy in stroke prediction, outperforming algorithms like Naive Bayes, Logistic Regression, SVM, and Decision Tree. The authors also used Ensembled Naive Bayes and Ensembled Decision Tree. The study contributes to developing an integrated learning model and refining the fixed structure of the algorithm. The authors in [22] evaluate ten machine learning models for stroke prediction, including Naive Bayes variants, Gradient Boosting, KNN, SVM, Decision Tree, Random Forest, Logistic Regression, and MLP. The study highlights the importance of early stroke diagnosis, and the KNN algorithm reaches the highest accuracy of 94%.

In [23], deep learning models are used to predict major adverse cerebrovascular events (MACE) after acute ischemic stroke (AIS). Using clinical and imaging data, models like DeepSurv and Deep-Survival Machines surpassed traditional survival methods. The validation results, including AUC, sensitivity, specificity, and other metrics, showcased the reliability of the models for personalized patient outcome predictions, helping clinical decision-making. The reviewed studies, summarized in Table 1, highlight diverse ML approaches and their substantial contributions to stroke prediction. These advances emphasize the potential of ML to improve stroke risk assessment, enabling proactive interventions and improved patient outcomes.

## III. Materials and Methods

In this study, the SMOTEEN [25] data balance technique is used to address the problem of data imbalance and PCA to select the relevant features. Five baseline ML classifiers (RF, KNN, SVM, XGB, and LR) are used for classification. In this paper four different optimization techniques are used for comparison, i.e. PSO, DE, BO, and ACO optimization. To further enhance the prediction of stroke, we have used three different Optimized Ensemble Approaches(Weighted, Soft Voting, and Stacked Ensemble) to find the most efficient solution. Figure 1 represents the stroke detection workflow diagram. and Algorithm 1 presents the data pre-processing and PCA-based feature selection.

**Fig. 1.**
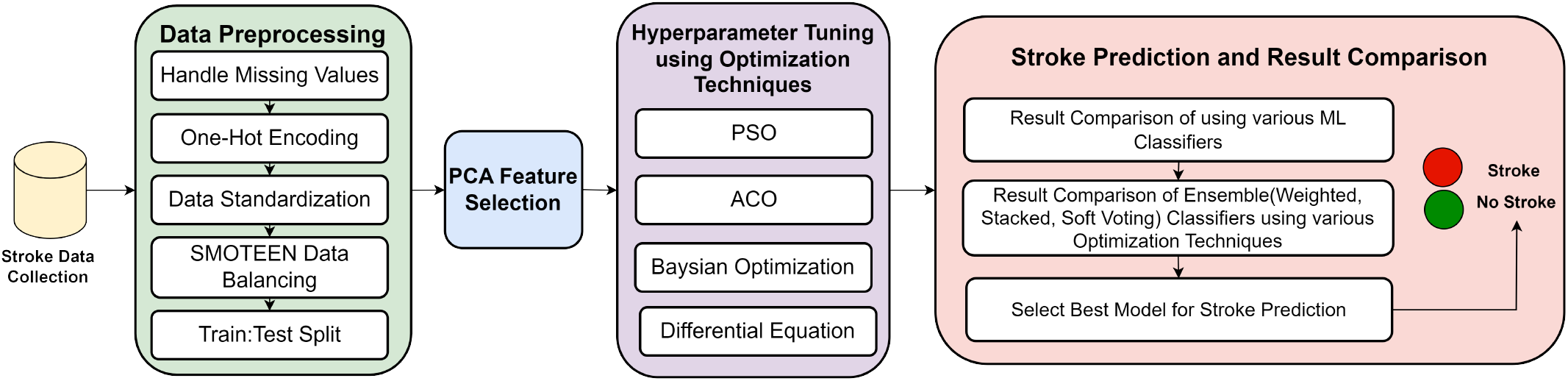
Stroke Prediction Workflow.

### A. Dataset

In this paper, we have used the Kaggle data set [24] that includes 5,110 patients and includes variables such as sex, age, hypertension status, history of heart disease, marital status, type of occupation, type of rehabilitation, average glucose level, body mass index (BMI), smoking behavior, and incidence of stroke. The main variable of interest is the prediction of the stroke, represented as a binary result (1 indicating the stroke, 0 indicating that there is no stroke). In this paper, SMOTEEN Data Balancing is used to address data imbalances to improve the accuracy of the model. Most of the samples obtained through various sampling methods were inadequate to provide adequate training data because class 0 had fewer instances, resulting in inaccurate results. Data preprocessing involves removing redundant values, selecting key features, and analyzing the data. A comparison of the classes and data points is illustrated in Figure 2 (classes before SMOTEEN) and Figure 3 (classes after SMOTEEN).

**Fig. 2.**
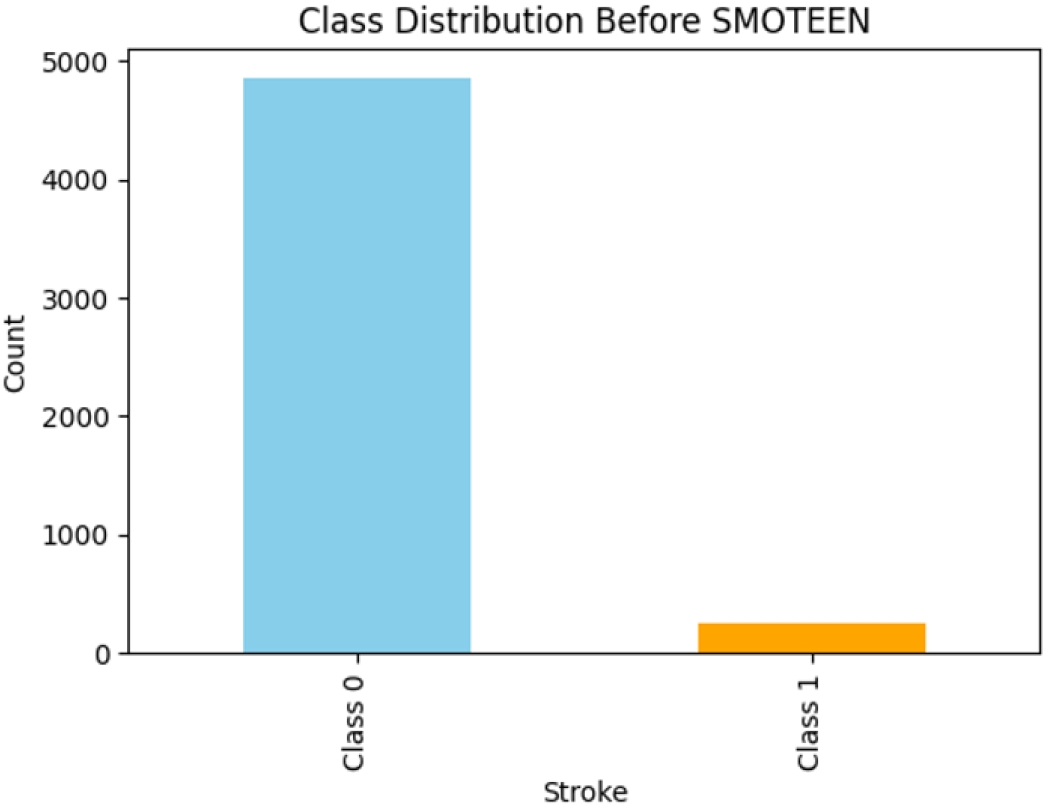
Class Distribution before Data Balancing.

**Fig. 3.**
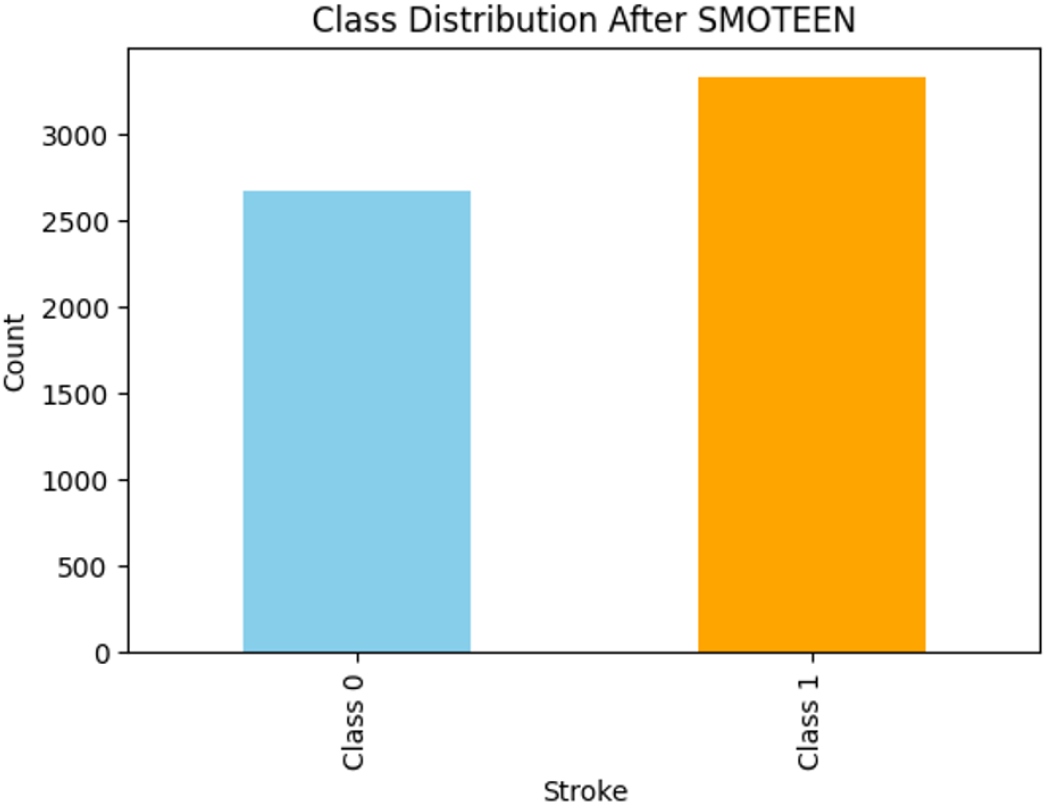
Class Distribution after SMOTEEN Data Balancing.

#### Algorithm 1 Data Preprocessing with SMOTEEN Balanching and PCA Feature Selection

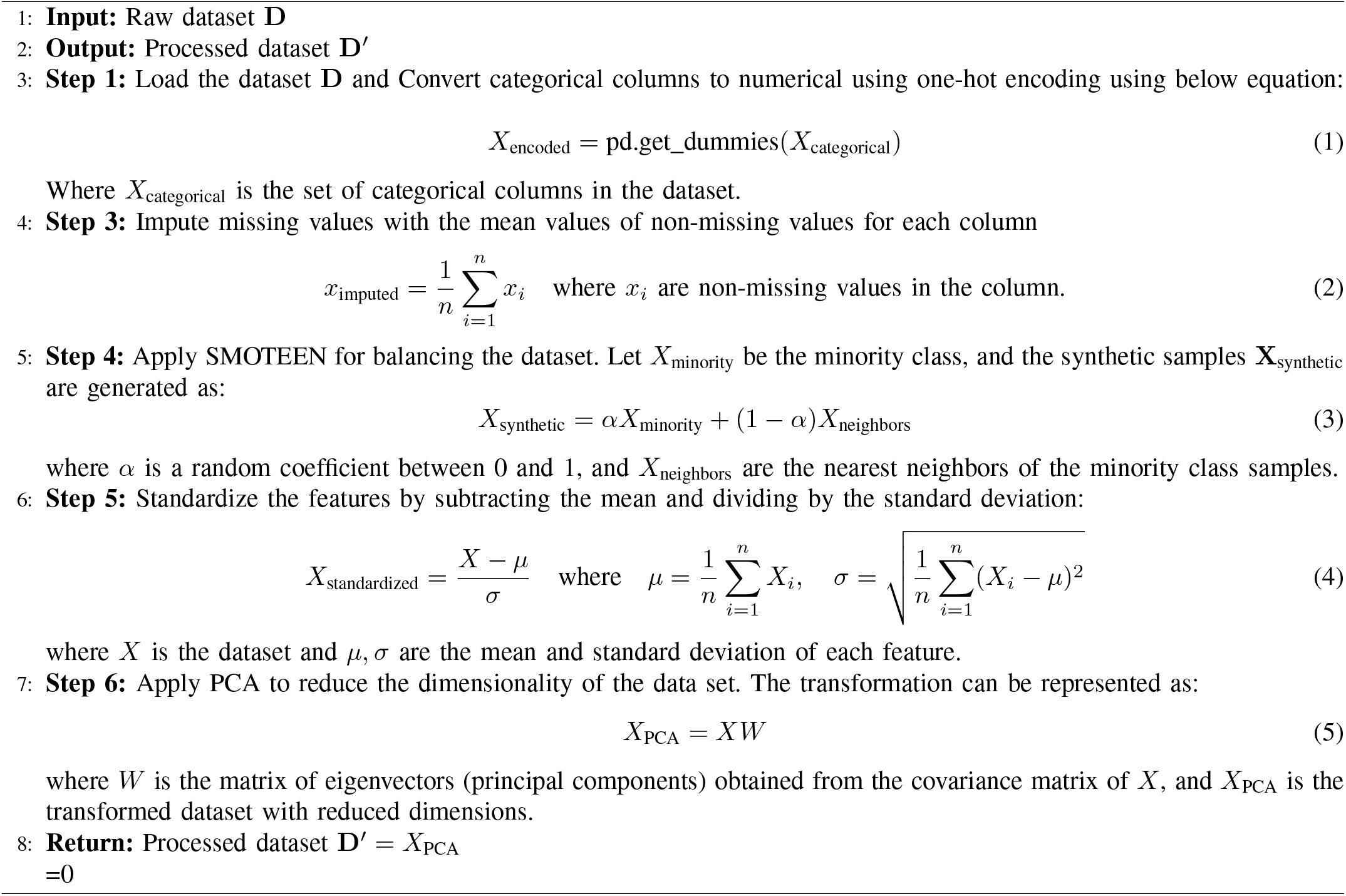

### B. PCA for Feature Selection

Principal Component Analysis (PCA) [26] is a method used for linear dimensionality reduction, converting data into a new coordinate system where the largest variances by any data projection are situated on the initial coordinates, known as principal components. The principal components are determined through eigenvalue decomposition of the data’s covariance matrix. Given a data matrix *X* with *n* samples and *p* characteristics, the covariance matrix Σ is calculated as:

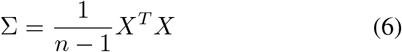

The next step is to perform the decomposition of the eigenvalues in the covariance matrix Σ, obtaining the eigen-values *λ*_1_, *λ*_2_, …, *λ*_*p*_ and the corresponding eigenvectors *v*_1_, *v*_2_, …, *v*_*p*_, such that:

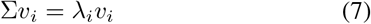

where *λ*_*i*_ represents the eigenvalue and *v*_*i*_ the corresponding eigenvector. The eigenvectors (principal components) are sorted in descending order based on their eigenvalues, and the first *k* eigenvectors form the new basis for the reduced feature space.

The data is then projected onto the new basis by multiplying the original data *X* by the matrix of the top *k* eigenvectors:

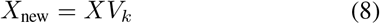

where *V*_*k*_ is the matrix of the top *k* eigenvectors, and *X*_new_ is the transformed data in the new *k* dimensional space. PCA is widely used to reduce the dimensionality of datasets while preserving as much variance as possible.

### C. Classifier Ensemble

*Stacked Ensemble Classifier(S*_*E*_): Stacking [27], or stacked generalization, is a hierarchical ensemble methodology that synthesizes multiple base learners into a meta-learner as shown in Figure 4. Formally, let the output of the base models *n* be represented as *Z* = *{P*_1_, *P*_2_, …, *P*_*n*_*}*. The metamodel *f*_meta_ learns a mapping *f*_meta_(*Z*) *→ y*, where *y* is the target variable.

**Fig. 4.**
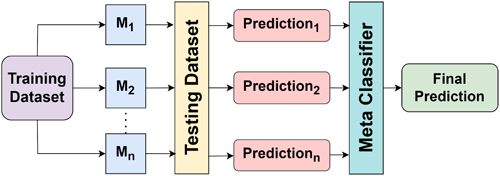
Stacked Ensemble Classifier Workflow.

The training process for stacked ensembles typically involves:

- Training base models independently on the dataset (*X, y*).
- Generating predictions (or transformed feature representations) from these models on either a holdout validation set or via cross-validation.
- Training a meta-model using the outputs of base models as input features and the true labels *y* as targets.

Depending on the complexity of the problem and the variety of base learners, the meta-model can utilize either linear or nonlinear mappings, including methods like logistic regression, gradient boosting, or neural networks. Stacking naturally harnesses the unique advantages of each model, typically achieving better generalization results. The effectiveness of the ensemble is influenced by the diversity among base models as well as the learning capability of the meta-model.

#### 2) Weighted Ensemble Classifier(W_E_)

The weighted ensemble method [28] is formulated to aggregate predictions from multiple base models by assigning a weight *w*_*i*_ to the output of each model based on its individual performance or importance as shown in Figure 5. The combined prediction *P*_ensemble_ is expressed mathematically as:

**Fig. 5.**
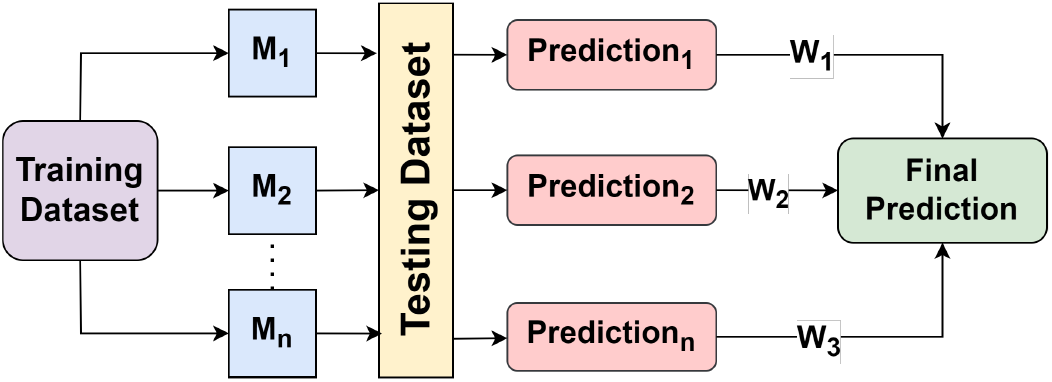
Weighted Ensemble Classifier Workflow.

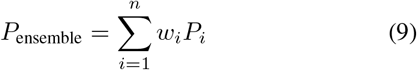

where *P*_*i*_ denotes the output of the *i*-th base model, and *w*_*i*_ represents its corresponding weight such that

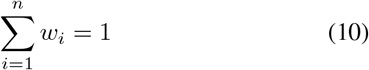

This method successfully tackles model variance by highlighting the input of models that perform well in predictions, all while preserving diversity. Nonetheless, it requires a strong approach to assigning weights to avoid dependency on one specific model, thereby maintaining the ensemble’s stability.

#### 3) Soft Voting Ensemble Classifier(SV_E_)

The voting ensemble method [29] consolidates predictions from multiple base models using either majority voting (hard voting) or probability averaging (soft voting) as shown in Figure 6. soft voting averages the probability distributions predicted by the models using the following formula:

**Fig. 6.**
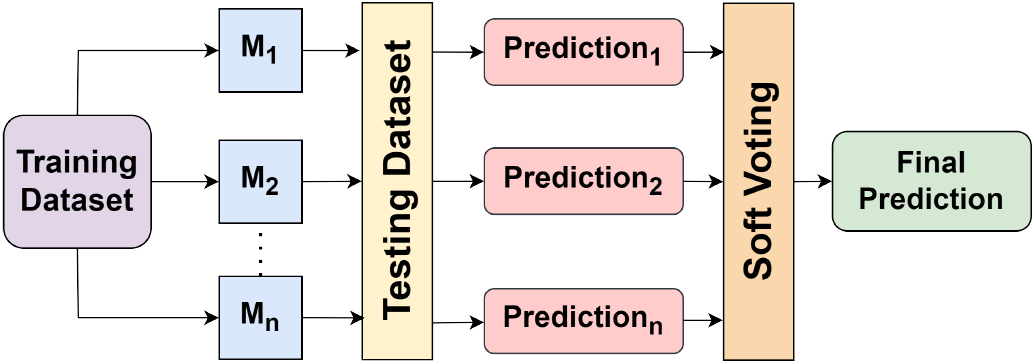
Soft Voting Ensemble Classifier Workflow.

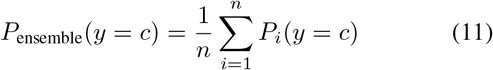

where *P*_*i*_(*y* = *c*) denotes the predicted probability of class *c* by model *i*. Voting ensembles are based on the assumption that base models are diverse and uncorrelated, since aggregate prediction smooths out individual model errors. This method avoids model weighting, making it computationally efficient and easy to implement.

### D. Optimization Techniques

#### 1) Differential Equation(DE)

Differential Evolution (DE) [30] is a strategy for optimization that is based on a population of potential solutions, which are refined through a cycle of mutation, crossover, and selection processes. During each cycle, a fresh candidate solution is produced by adding a scaled disparity between two population members with a third one. The fundamental mutation equation in DE is expressed as:

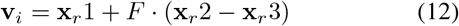

where **v**_*i*_ is the mutated vector for the *i*-th individual, **x**_*r*_1, **x**_*r*_2, **x**_*r*_3 are randomly selected population vectors, and *F* is a scaling factor. The crossover operation creates a trial vector **u**_*i*_ from the parent vector **x**_*i*_ and the mutated vector **v**_*i*_:

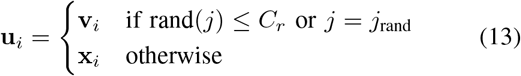

where *C*_*r*_ is the crossover rate, rand(*j*) is a random value between 0 and 1, and *j*_rand_ is a randomly selected index. The best candidate solution is selected based on the objective function evaluation, ensuring convergence towards an optimal solution.

#### 2) Bayseian Optimization(BO)

Bayesian Optimization (BO) [31] is a technique to optimize costly-to-evaluate objective functions using a probabilistic model. It creates a surrogate probabilistic model, often employing a Gaussian process (GP), to represent the objective function. An acquisition function is then utilized to direct the search towards the optimal solution. The GP model is described by:

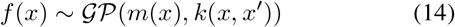

where *f* (*x*) is the objective function, *m*(*x*) is the mean function and *k*(*x, x*^*′*^) is the covariance function (kernel). The acquisition function, which is used to decide the next evaluation point, is typically based on the expected improvement (EI). The expected improvement is given by:

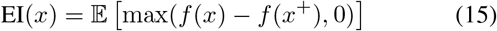

where *x*^+^ is the best observed point in the current, and *f* (*x*) is the predicted value of the objective function at the point *x*. The next point to evaluate is chosen by maximizing the acquisition function.

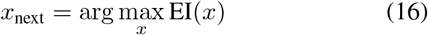

By iterating this process, BO efficiently finds the global optimum while minimizing the number of evaluations of objective functions.

#### 3) Particle Swarm Optimization(PSO)

Particle Swarm Optimization (PSO) [32] is a population-based optimization algorithm inspired by the collective behavior of bird flocks. In this approach, each particle symbolizes a possible solution, continuously refining its position by considering its own optimal past position as well as the optimal positions discovered by the swarm as a whole. The updates for position and velocity in PSO are defined as follows:

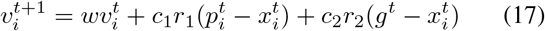

where 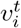 is the velocity of the *i*-th particle at iteration *t, w* is the inertia weight, *c*_1_ and *c*_2_ are acceleration coefficients, *r*_1_ and *r*_2_ are random values between 0 and 1, 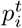 is the best personal position of the particle *i* -th particle, and *g*^*t*^ is the best global position of the swarm.

The position of the *i*-th particle is updated as:

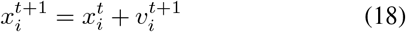

where 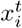 is the position of the *i*-th particle at iteration *t*. The particles move through the solution space by adjusting their velocities based on their own best experiences and the best global experience found by the swarm. This process continues until convergence or a stopping criterion is met.

#### 4) Ant Colony Optimization (ACO)

ACO is a natureinspired optimization algorithm based on the foraging behavior of ants [33]. The algorithm simulates how ants deposit pheromones on paths and adjust their movements to find the shortest path to a food source. ACO is used for discrete optimization problems, such as the traveling salesman problem. The core principle of ACO is to iteratively improve solutions by strengthening paths that lead to better solutions, using a pheromone updating rule.

The pheromone update formula is given by:

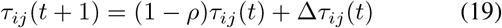

where: - *τ*_*ij*_(*t*) is the pheromone level on edge (*i, j*) at time *t*, - *ρ* is the evaporation rate, - Δ*τ*_*ij*_(*t*) is the pheromone deposited by ants on edge (*i, j*).

The probability of an ant choosing the next node in the solution is given by:

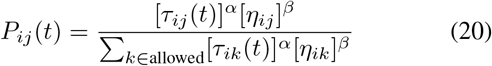

where: 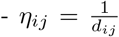is the inverse of the distance between nodes *i* and *j*, - *α* and *β* control the influence of pheromone and distance, respectively.

### E. Machine Learning Classifiers

#### 1) Logistic Regression(LR)

Logistic regression (LR) is a statistical method used for binary classification tasks. Models the probability of an outcome using a logistic function, ensuring that the output ranges between 0 and 1. The hypothesis for LR is given by:

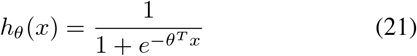

where *h*_*θ*_(*x*) represents the predicted probability, *θ* is the parameter vector and *x* is the input feature vector. The model is trained by minimizing the log-loss function, which measures the difference between predicted and actual labels, ensuring robust performance on classification tasks.

#### 2) Random Forest

Random Forest (RF) is an ensemble learning method that is used primarily for classification and regression tasks. It constructs multiple decision trees during training and outputs the class label based on majority voting or averages the predictions for regression. The prediction of the Random Forest classifier can be expressed as:

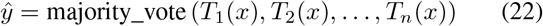

where *T*_*i*_(*x*) represents the prediction of the *i*-th decision tree for input *x*, and *n* is the total number of trees. By aggregating predictions, RF reduces overfitting and improves model generalization, making it effective for handling high-dimensional and imbalanced datasets.

#### 3) K Nearest Neighbor

The K-Nearest Neighbors (KNN) algorithm is a simple and intuitive supervised learning method that is used for classification and regression tasks. It works by identifying the *k* closest data points to a query point in the feature space based on a chosen distance metric, such as the Euclidean distance. The Euclidean distance *d* between two points **x**_**1**_ = (*x*_11_, *x*_12_, …, *x*_1*n*_) and **x**_**2**_ = (*x*_21_, *x*_22_, …, *x*_2*n*_) is calculated as:

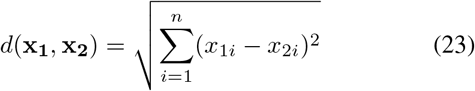

For classification, KNN assigns the query point to the class that is most frequent among its *k*-nearest neighbors. For regression, the output is typically the mean or median of the neighbors’ values. KNN is non-parametric and requires no explicit training phase, making it straightforward to implement but computationally intensive for large datasets.

#### 4) Extreme Gradient Boosting

Extreme Gradient Boosting(XGBoost) is an advanced implementation of gradient boosting, designed for speed and performance in both classification and regression tasks. It combines the predictions of an ensemble of weak learners, typically decision trees, by minimizing a differentiable loss function by using gradient descent. The model is built iteratively and each tree corrects the errors of the previous ones.

The objective function of XGBoost is given as:

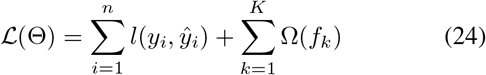

where:

- *l*(*y*_*i*_, *ŷ*_*i*_) is the loss function that measures the difference between the actual value *y*_*i*_ and the predicted value *ŷ*_*i*_.
- Ω(*f*_*k*_) is the regularization term to penalize the complexity of the *k*-th tree *f*_*k*_, expressed as:

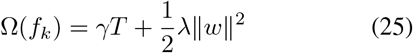

where *T* is the number of leaves in the tree, *γ* is the penalty for adding a leaf, *λ* controls the regularization of the weights of the *L*_2_ leaves *w*.

XGBoost employs the second-order Taylor expansion to approximate the loss function, improving optimization efficiency. Its ability to handle missing data, regularization, and parallel processing makes it highly effective for structured data tasks.

#### 5) Support Vector Machine

Support Vector Machine (SVM) is a supervised learning algorithm used for classification and regression tasks. It seeks to find the optimal hyperplane that maximizes the margin between data points of different classes. The hyperplane is defined as:

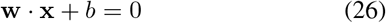

where **w** is the weight vector, **x** is the feature vector and *b* is the bias. For a correctly classified point (**x**_*i*_, *y*_*i*_), the condition is:

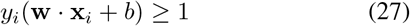

The objective of SVM is to minimize the norm *∥***w***∥* ^2^ subject to the above constraint, which can be formulated as:

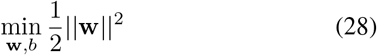

For nonlinearly separable data, a kernel function *K*(**x**_*i*_, **x**_*j*_) is used to map the input data into a higher-dimensional space to find a separating hyperplane.

## IV. Proposed System Architecture

The proposed system aims to predict stroke by applying the PCA feature reduction technique and the optimized ensemble classification approach. We have first applied the SMOTEEN Data Balancing Approach to balance the dataset and after that Five Baseline ML Classifiers are used. Then, to further enhance model performance, optimization techniques such as Ant Colony Optimization (ACO), Differential Equation(DE), Bayesian Optimization(BO), and Ant Colony Optimization(ACO) are applied to perform optimized ensemble methods (stacked, soft voting, and weighted average). The ensemble model, using optimized parameters, combines the predictions from baseline ML models to improve accuracy.Accuracy, precision, recall, F1 score, and AUC, with visualizations have been used for performance evaluation.ROC curves and confusion matrices are used to provide a comprehensive evaluation of the model’s predictive capabilities. This approach ensures robust stroke prediction while addressing class imbalance, making the model more reliable and accurate. Figure 7 and Algorithm 2 show the complete proposed system architecture.

**Fig. 7.**
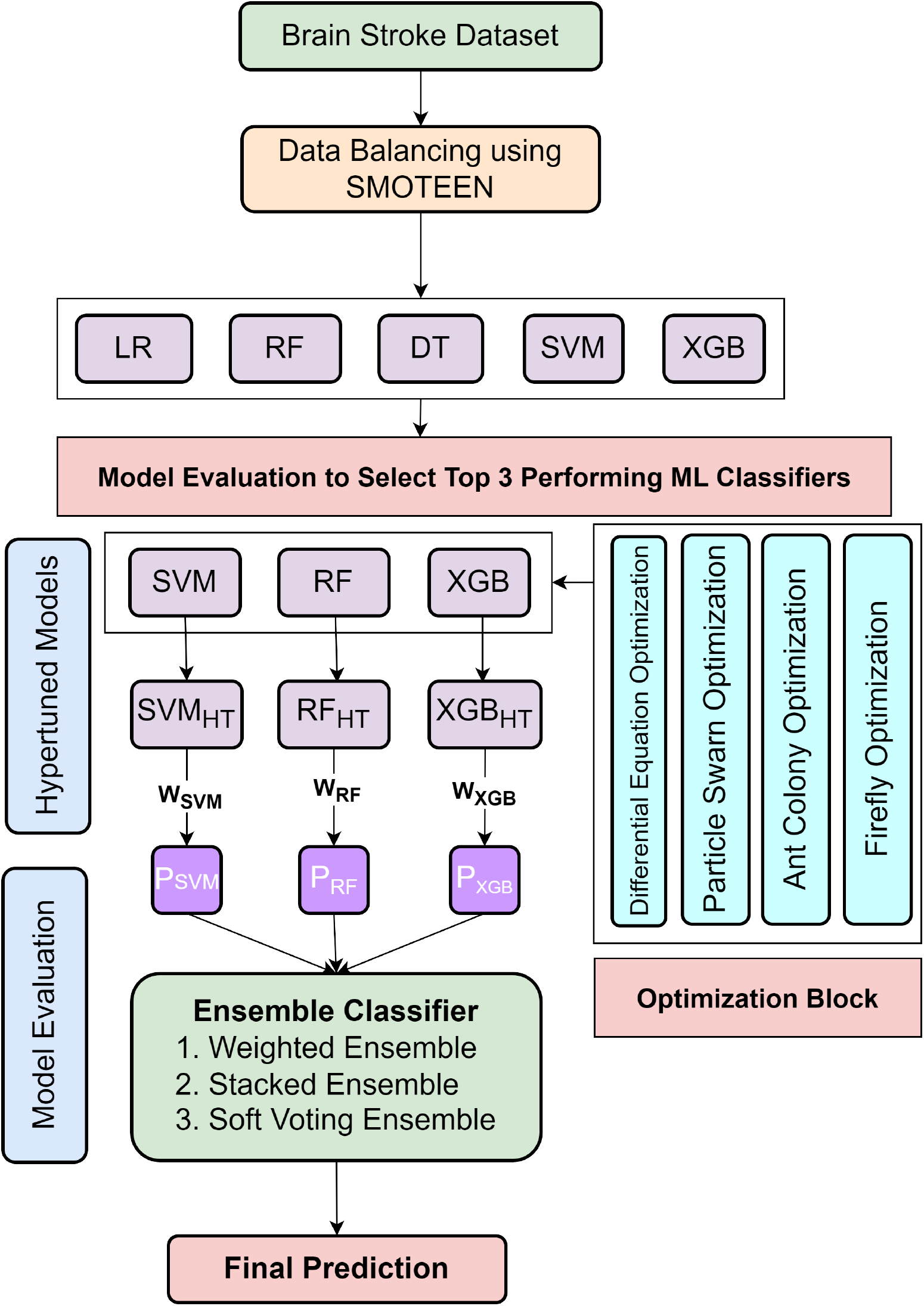
Proposed System Architecture.

## V. Experimental Setup

Stroke dataset is processed over proposed methodology. In this process, imbalance dataset problems are also addressed.In this work a highly imbalanced Kaggle brain stroke dataset with data preprocessing as explained in Algorithm 1. In this experiment, 80% of the data is used to train the ML classifiers, and the remaining 20% is used to test them, ensuring a robust evaluation. The empirical results of the baseline ML classifiers and optimized Ensemble ML classifiers on the SMOTEEN balanced data set have been represented using different visualization techniques based on the evaluation metrics considered, viz. Accuracy, precision, recall, F1 score, and AUC-ROC. We have implemented the experiments on Kaggle Platform with Python3, GPU Support and 30GB RAM. To evaluate the model performance, the following metrics are used, and in equations the following abbreviations are used: TP(True Positives), TN(True Negatives), FP(False Positives), FN(False Negatives).

1. **Accuracy(ACC)**:

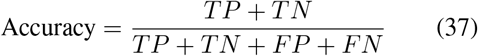
2. **Precision(P)**:

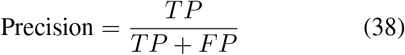
3. **Recall(R)**:

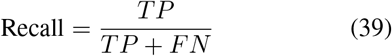
4. **F1 Score**

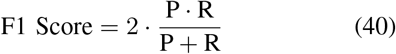
5. **ROC AUC (Receiver Operating Characteristic Area Under the Curve)**: This metric shows the performance of models based on specificity and sensitivity. The AUC (Area Under the Curve) value ranges from 0 to 1, where a value closer to 1 indicates a better-performing model.

## VI. Results and Discussion

### A. Results on Imbalanced Dataset

The table I summarizes the performance metrics of various machine learning models (LR, RF, KNN, SVM, and XGB) for two classes (0 and 1). Across all models, the metrics for Class 0 (No Stroke) are significantly better than those for Class 1 (Stroke), highlighting a severe data imbalance problem as the number of class 1 instances is significantly lower than class 0. For Class 0, all models achieve high precision, recall, and F1 scores, with accuracies ranging from 0.936 to 0.942. However, for Class 1, the metrics are notably poor, with precision as low as 0.25, recall close to 0, and F1-scores below 0.08 for most models. The Area Under the Precision-Recall Curve (AUCPR) values are also low, indicating that the models struggle to distinguish the minority class. This disparity underscores the challenge of imbalanced datasets, where models are biased toward the majority class, leading to poor performance in the minority class. Figures 8 and Figure 9 show the comparative ROC and the confusion matrix of the baseline ML classifications.

**TABLE I.**
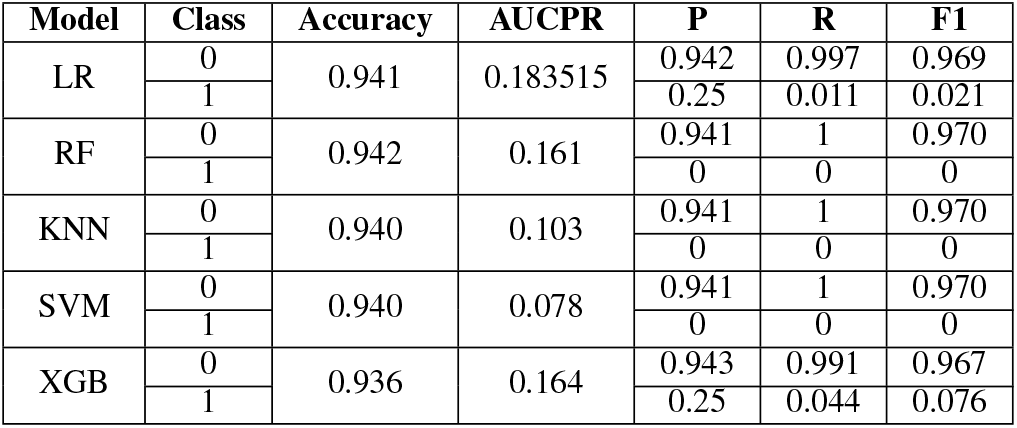
Performance metrics for different models.

| Model | Class | Accuracy | AUCPR | P | R | F1 |
| --- | --- | --- | --- | --- | --- | --- |
| LR | 0 | 0.941 | 0.183515 | 0.942 | 0.997 | 0.969 |
|  | 1 |  |  | 0.25 | 0.011 | 0.021 |
| RF | 0 | 0.942 | 0.161 | 0.941 | 1 | 0.970 |
|  | 1 |  |  | 0 | 0 | 0 |
| KNN | 0 | 0.940 | 0.103 | 0.941 | 1 | 0.970 |
|  | 1 |  |  | 0 | 0 | 0 |
| SVM | 0 | 0.940 | 0.078 | 0.941 | 1 | 0.970 |
|  | 1 |  |  | 0 | 0 | 0 |
| XGB | 0 | 0.936 | 0.164 | 0.943 | 0.991 | 0.967 |
|  | 1 |  |  | 0.25 | 0.044 | 0.076 |

**Fig. 8.**
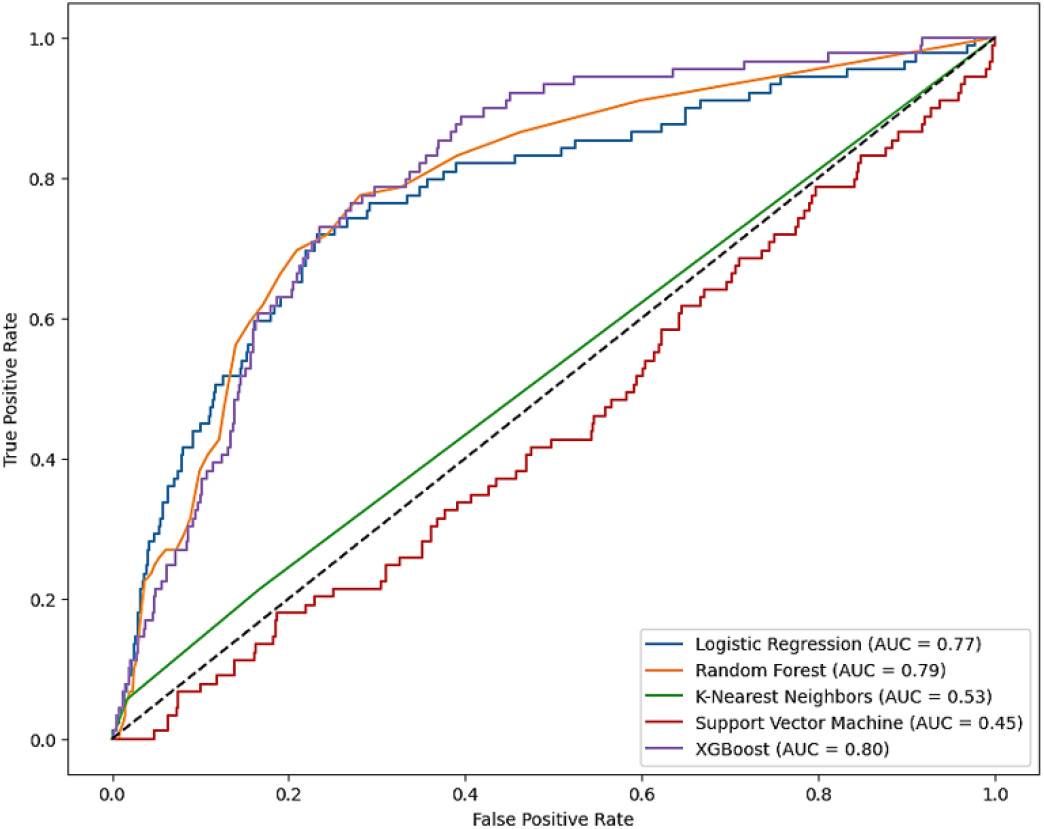
ROC Comparison for Imbalanced Dataset.

**Fig. 9.**
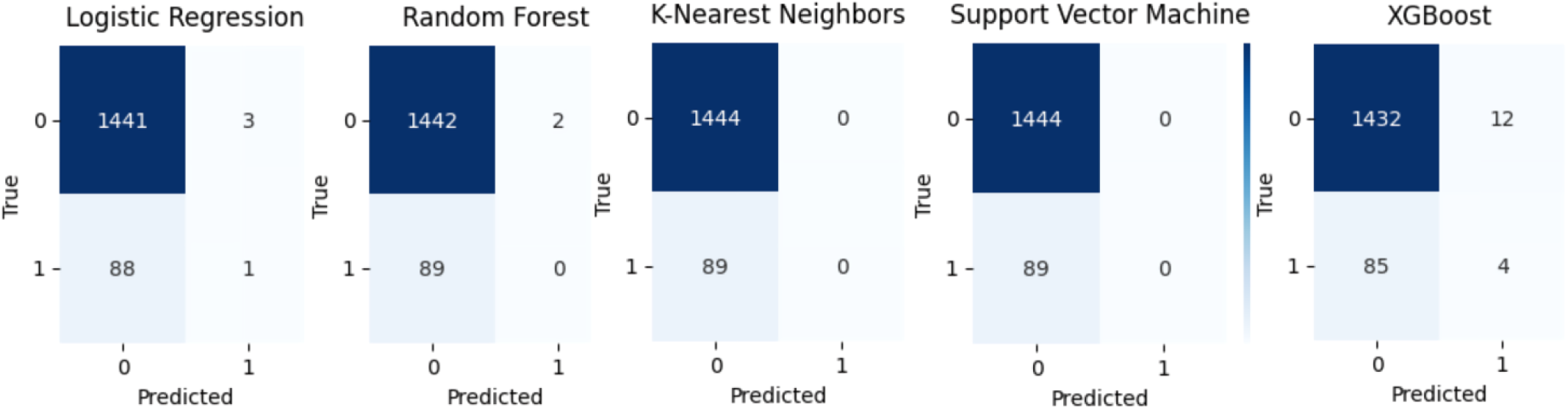
Confusion Matrix for Imbalanced Dataset.

### B. Results on Balanced Dataset using SMOTEEN and PCA Feature Selection

The table II shows that all models perform well, achieving high precision, recall, and F1 scores for both the majority (Class 0) and minority (Class 1) classes. Random Forest achieves the highest accuracy of 94.4% and an AUCPR of 0.993, with a strong performance in handling the minority class, achieving a precision of 0.98, recall of 0.90, and an F1 score of 0.94. XGB and SVM also perform exceptionally well, with accuracies of 94.2% and 91.7%, respectively. Based on these results, the top three classifiers: Random Forest, XGBoost, and SVM were selected for further optimization using an ensemble approach. Figures 10 and Figure 11 show the comparative ROC and the confusion matrix of the baseline ML classifications.

**TABLE II.**
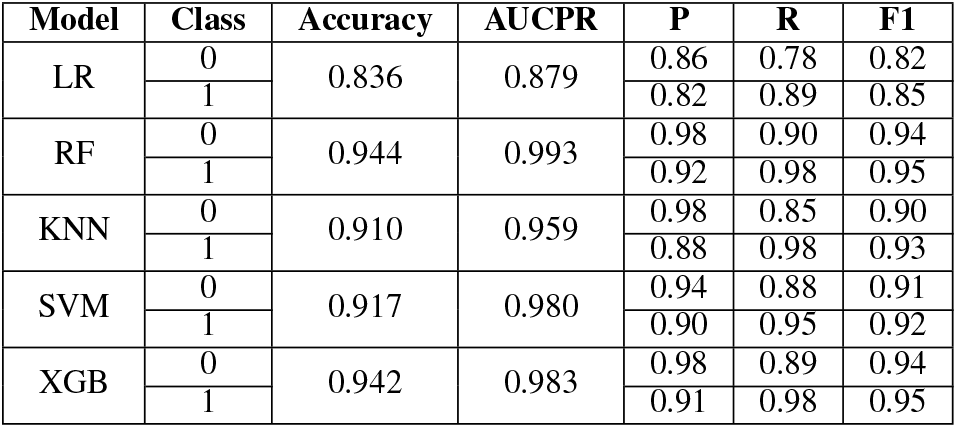
Performance metrics for different models.

**Fig. 10.**
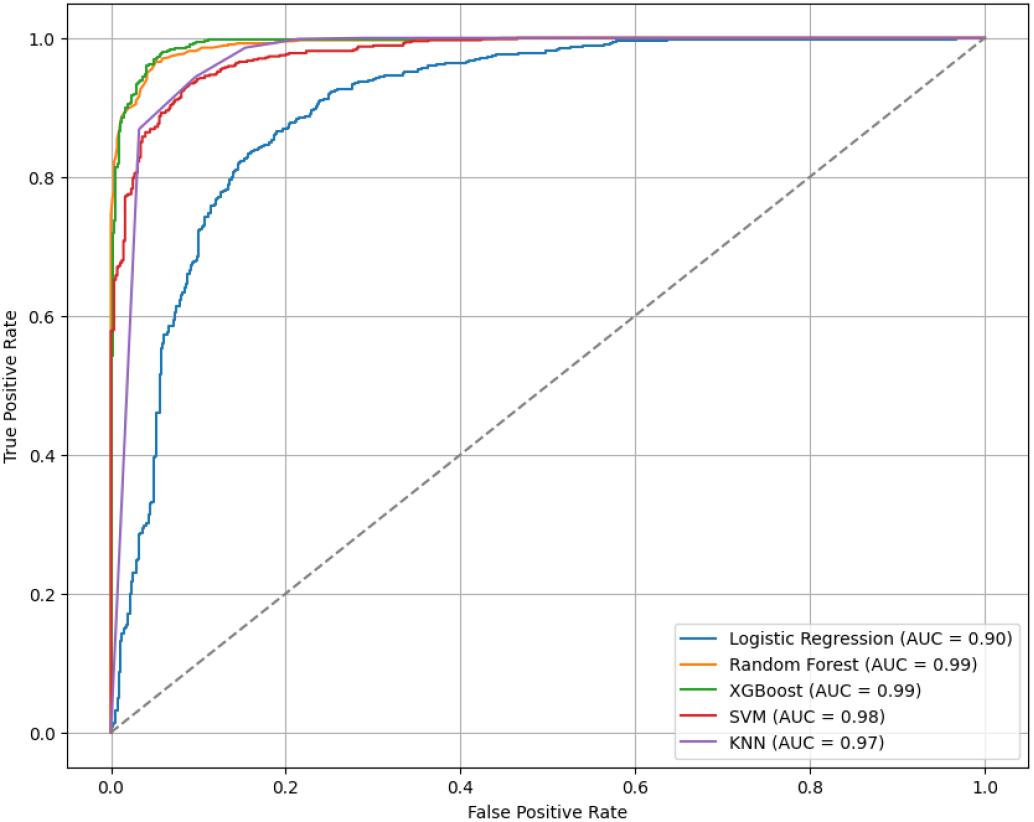
ROC Comparison for SMOTEEN Balanced Dataset with PCA Feature Selection.

**Fig. 11.**
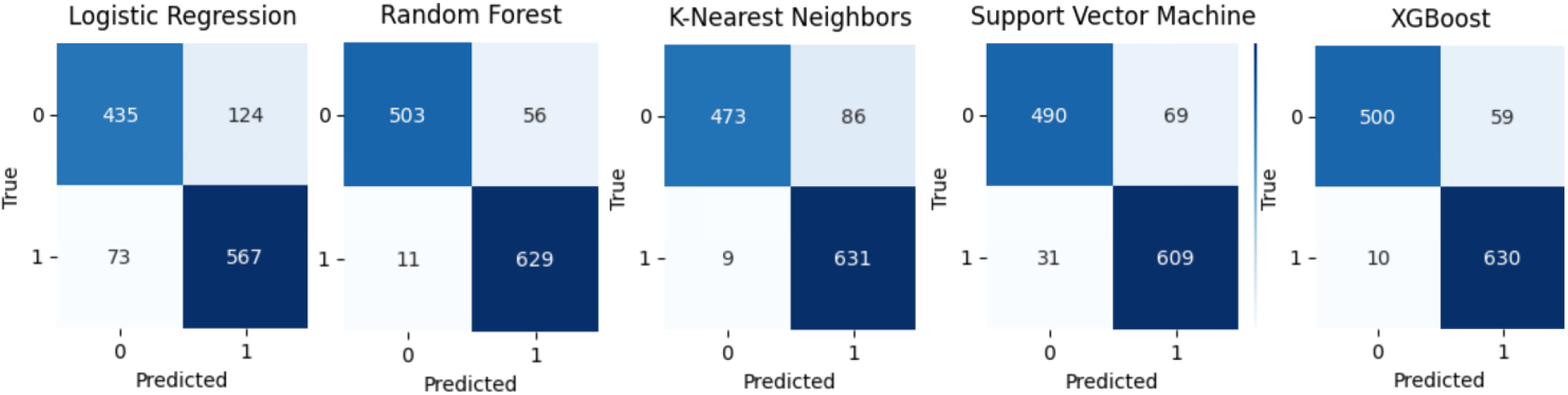
Confusion Matrix for SMOTEEN Balanced Dataset with PCA.

### C. Effect of Data Balancing

The results shown in Figure 12 highlight the significant improvement achieved by applying SMOTEEN to address the class imbalance in the dataset. For Class 1 (Stroke), the imbalanced dataset shows very poor performance, with mean precision, recall, and F1-score values of 0.1296, 0.0378, and 0.0476, respectively. However, after SMOTEEN application, these metrics improve substantially to 0.886, 0.956, and 0.920,\ respectively. This improvement is visually evident in the figure, where the balanced dataset using SMOTEEN outperforms the imbalanced dataset with significant scores. These findings demonstrate SMOTEEN’s ability to effectively generate synthetic minority class samples while reducing noise, thereby enhancing the classifier’s ability to detect stroke cases and deliver a more balanced performance.

**Fig. 12.**
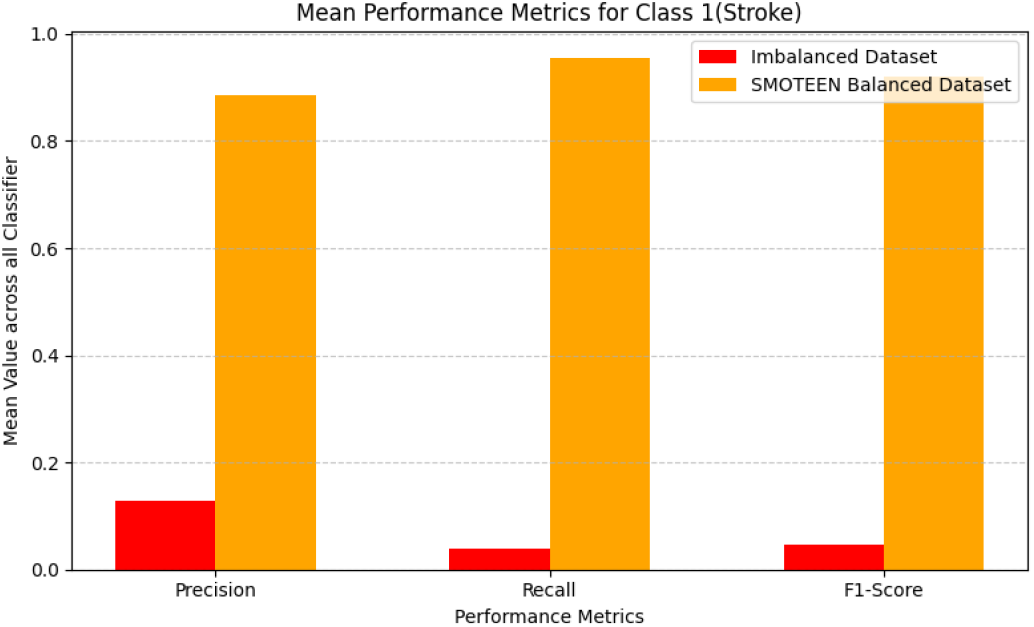
Mean Performance Metric Comparison between Imbalanced and SMOTEEN Balanced Dataset.

#### Algorithm 2 Proposed System Workflow Steps

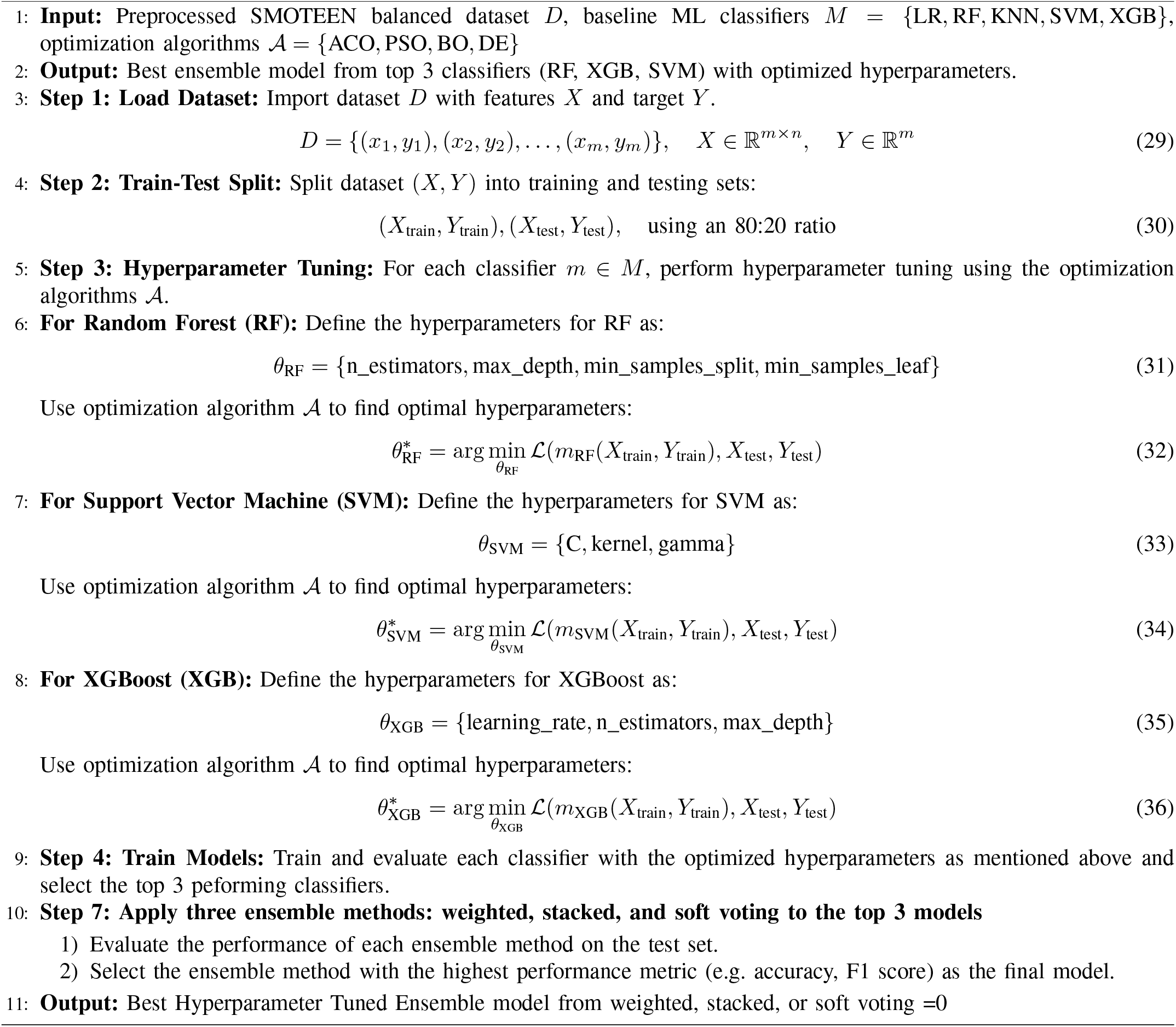

Figure 13 highlights the significant impact of data balance using the SMOTEEN technique on the AUCPR values of various models. In the imbalanced dataset, models show poor performance in handling the minority class, with lower AUCPR values, especially for Logistic Regression with an AUCPR of 0.183 and SVM with an AUCPR of 0.078. After applying SMOTEEN, the AUCPR values improve considerably for all models, with Random Forest achieving 0.993 and XGB reaching 0.983. This demonstrates that data balancing effectively reduces bias towards the majority class and enhances the ability of models to classify minority instances accurately.

**Fig. 13.**
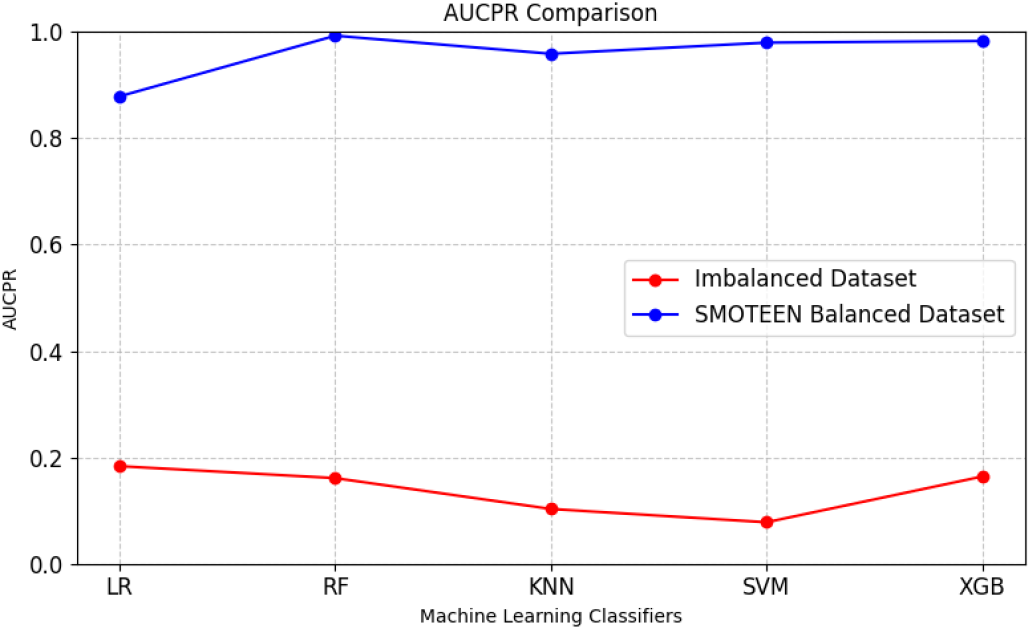
AUCPR Comparison between Imbalanced and SMOTEEN Balanced Dataset.

### D. Results on Proposed Model

Table III presents a comparison of the performance of various optimization techniques, including ACO, DE, BO, and PSO, applied to different models: RF, SVC, GB, *W*_*E*_, *S*_*E*_, *SV*_*E*_. Among all the approaches, the PSO-*S*_*E*_ combination achieves the best results, with the highest accuracy of 0.956, precision of 0.956, recall of 0.956, F1 score of 0.992, and AUC of 0.993, highlighting its superior ability to balance classification performance and effectively handle imbalanced data. In contrast, the ACO-XGB combination performs the poorest, with the lowest accuracy of 0.897, precision of 0.878, and AUC of 0.895, indicating weak generalization and classification capabilities. Overall, PSO proves to be the most effective optimization technique, consistently delivering high performance across models, whereas ACO shows significant under performance, particularly with the GB model. The comparison of the confusion matrix for all 12 different optimized ensemble models is shown in Figure 14.

**TABLE III.** Performance Metrics for Different Optimization Techniques.

| Optimization | Model | Accuracy | Precision | Recall | F1-Score | AUC |
| --- | --- | --- | --- | --- | --- | --- |
| ACO | RF | 0.943 | 0.914 | **0.986** | 0.949 | 0.992 |
|  | SVC | 0.922 | 0.914 | 0.942 | 0.928 | 0.978 |
|  | GB | 0.897 | 0.878 | 0.938 | 0.907 | 0.895 |
| | $W_E$ | 0.941 | 0.913 | 0.983 | 0.947 | 0.987 |
| | $S_E$ | 0.945 | 0.921 | 0.981 | 0.950 | 0.987 |
| | $SV_E$ | 0.951 | 0.940 | 0.970 | 0.955 | 0.992 |
| DE | RF | 0.938 | 0.943 | 0.935 | 0.937 | 0.988 |
|  | GB | 0.930 | 0.933 | 0.927 | 0.929 | 0.979 |
|  | SVC | 0.937 | 0.938 | 0.935 | 0.936 | 0.987 |
| | $W_E$ | 0.948 | 0.952 | 0.945 | 0.947 | 0.990 |
| | $S_E$ | 0.945 | 0.948 | 0.942 | 0.944 | 0.989 |
| | $SV_E$ | 0.946 | 0.949 | 0.944 | 0.945 | 0.990 |
| BO | RF | 0.942 | 0.947 | 0.939 | 0.941 | 0.989 |
|  | GB | 0.934 | 0.937 | 0.932 | 0.934 | 0.980 |
|  | SVC | 0.937 | 0.938 | 0.935 | 0.936 | 0.987 |
| | $W_E$ | 0.949 | 0.951 | 0.949 | 0.949 | 0.780 |
| | $S_E$ | 0.949 | 0.950 | 0.949 | 0.949 | 0.989 |
| | $SV_E$ | 0.947 | 0.948 | 0.947 | 0.947 | 0.990 |
| PSO | RF | 0.941 | 0.946 | 0.938 | 0.940 | 0.987 |
|  | GB | 0.923 | 0.925 | 0.920 | 0.922 | 0.972 |
|  | SVC | 0.936 | 0.937 | 0.934 | 0.935 | 0.987 |
| | $W_E$ | 0.953 | 0.954 | 0.953 | 0.953 | 0.992 |
| | $S_E$ | **0.956** | **0.956** | 0.956 | **0.992** | **0.993** |
| | $SV_E$ | 0.952 | 0.951 | 0.951 | 0.991 | 0.992 |

**Fig. 14.**
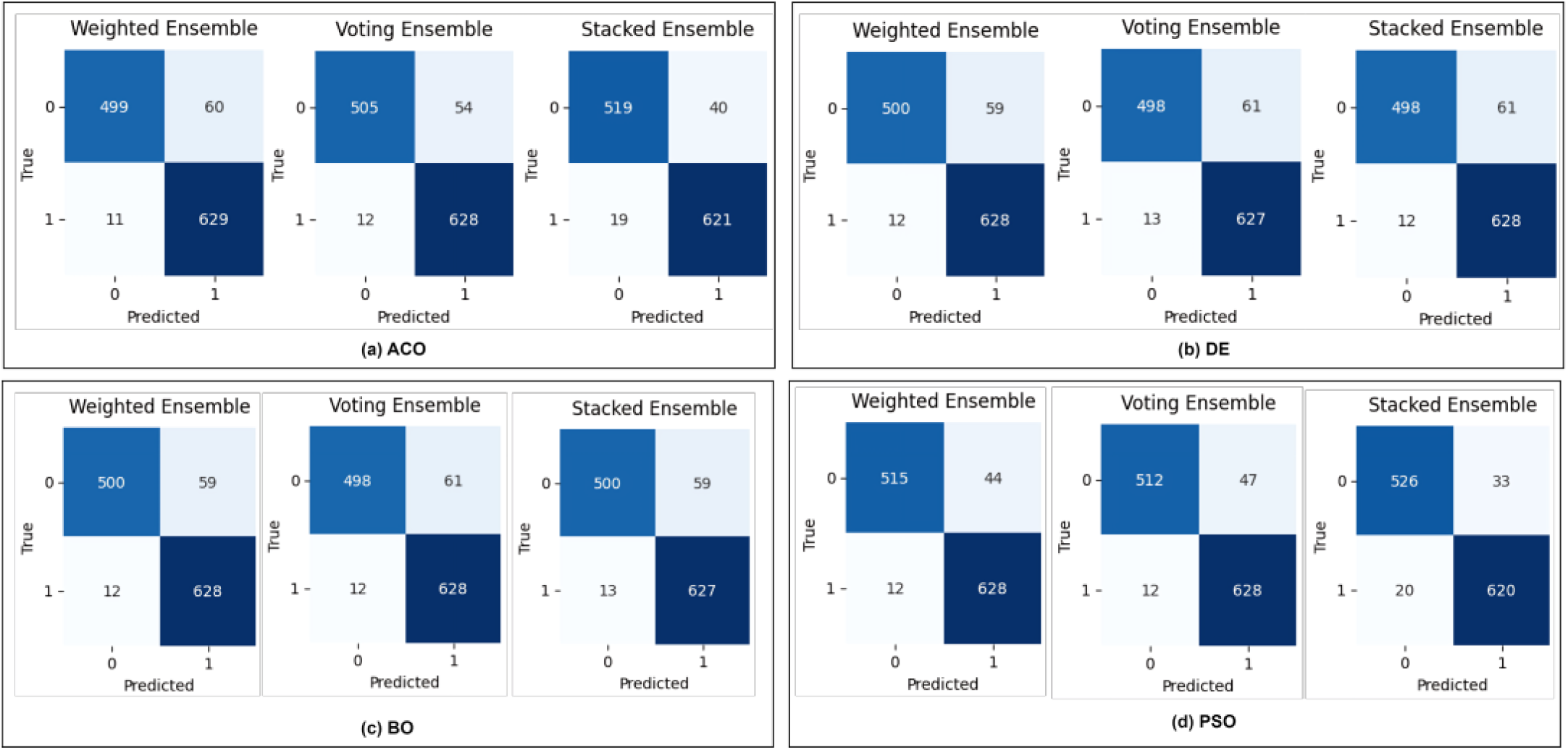
Confusion Matrix: (a) Ant Colony Optimization Based Ensemble Models, (b) Differential Equation Based Ensemble Models, (c) Bayesian Optimization based Ensemble Models, (d) Particle Swarm Optimization based Ensemble Models.

We have also verified our proposed(PSO Optimized Stacked Ensemble) Model performance against well established baseline ML Classifiers on SMOTEEN Balanced Dataset with PCA Feature selection technique. The results shown in Figure 15 showed that our proposed model consistently achieves the highest values, with an accuracy of 0.956, precision of 0.956, recall of 0.956 and F1-Score of 0.992, outperforming all other models. The proposed model achieves the highest F1 score of 0.992, significantly outperforming all other models. Compared to the second-best model, RF, which has an F1-Score of 0.945, the proposed model shows a notable improvement of 4.97%, demonstrating its superior ability to balance Precision and Recall. RF performs strongly, followed by KNN, SVM, and XGB, all of which achieve an F1 score of 0.915, indicating competitive but slightly lower performance. However, LR lags behind, with the lowest F1 score of 0.835, making it the weakest model in terms of balanced classification performance.

**Fig. 15.**
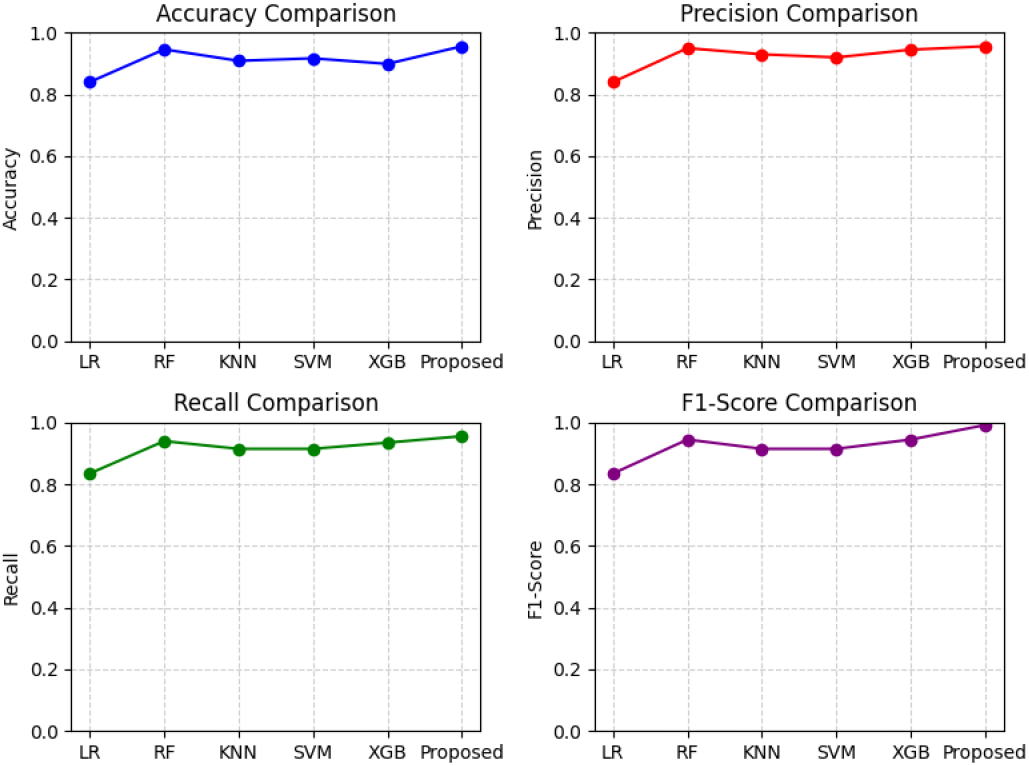
Evaluation Metrics Comparison of Baseline ML Classifiers with Proposed Model.

Table IV compares the results of the proposed approach with previous work done on the same data set. The results show that our proposed approach outperforms other existing work which shows the effectiveness of PSO Optimized Stacked Ensemble ML Classifier over baseline ML Classifiers.

**TABLE IV.**
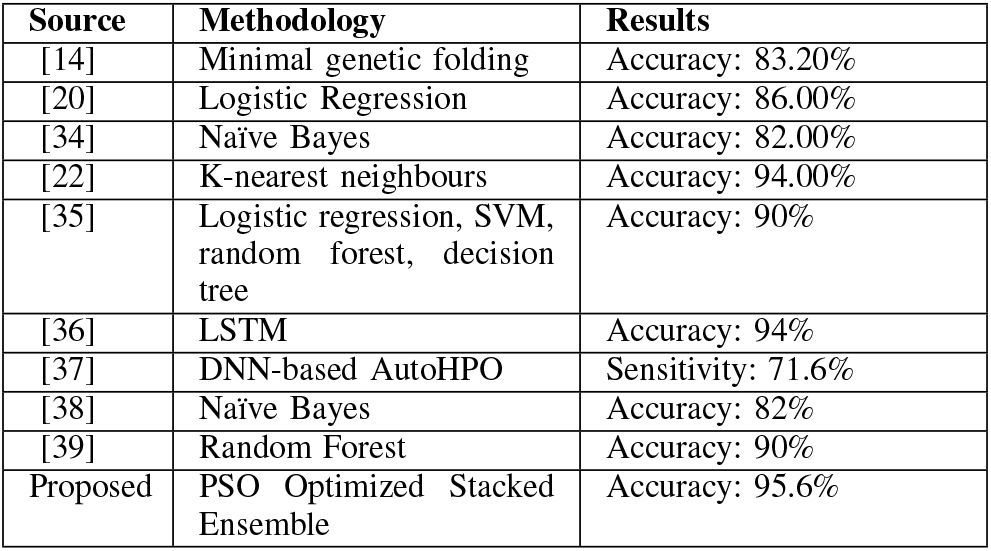
Comparison of Proposed Approach with Existing Work.

### E. Statistical Analysis

To evaluate the strength and robustness of the optimization techniques and different ensemble methods used in this paper, we carried out Statistical Analysis. In this we have compared the difference between Best Accuracy(*B*_*A*_) and Mean Accuracy(*M*_*A*_). The standard deviation (SD) is also calculated from the accuracy distribution for all proposed models. The formulas are shown in the following equations.

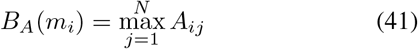

Where: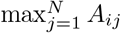: Maximum accuracy for model *m*_*i*_ across all models.

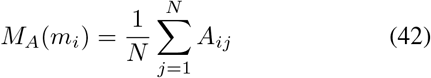

Where: *N* : Total number of ETL models, *A*_*ij*_: Accuracy of model *m*_*i*_ for ETL model *j, M*_*A*_(*m*_*i*_): Mean accuracy of model *m*_*i*_.

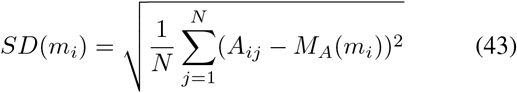

Where: *A*_*ij*_: Accuracy of model *m*_*i*_ for ETL model *j, M*_*A*_(*m*_*i*_): Mean accuracy of model *m*_*i*_, The sum of squared deviations is divided by *N* and the square root is taken to compute the standard deviation.

The table V presents the performance metrics of three ensemble models (*W*_*E*_, *S*_*E*_ and *SV*_*E*_) based on Mean Accuracy (*M*_*A*_), Mean Precision (*M*_*P*_), Mean Recall (*M*_*R*_), Mean F1 score (*M*_*F*_ 1), Best Accuracy (*B*_*A*_), the difference between Best Accuracy and Mean Accuracy (*B*_*A*_-*M*_*A*_), and Standard Deviation (STD). Among the models, SE achieves the highest *M*_*F*_ 1 (0.959) and *B*_*A*_ (0.956), indicating its superior ability to balance precision and recall while achieving the best classification performance. However, *SV*_*E*_ demonstrates the highest consistency, with the lowest *B*_*A*_-*M*_*A*_ (0.003) and STD (0.003), making it the most stable model. *W*_*E*_ performs slightly lower across all metrics, with an *M*_*F*_ 1 of 0.949 and *B*_*A*_ of 0.953, but still delivers competitive results. Overall, *S*_*E*_ is the best performing model in terms of overall accuracy and F1 score, while *SV*_*E*_ is the most reliable due to its minimal variability.

**TABLE V.**
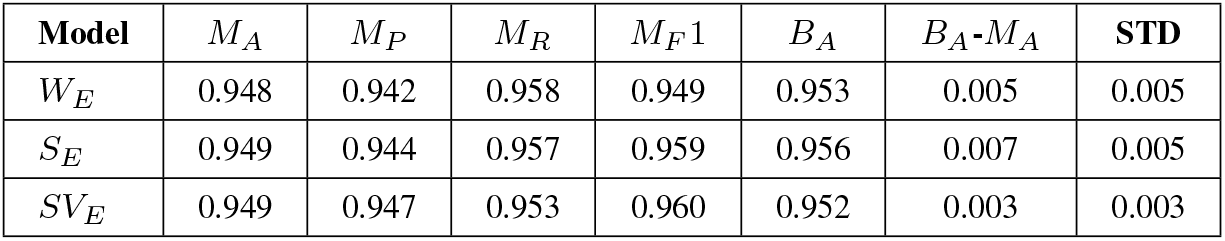
Statistical Analysis of Ensemble Models across all Optimizers.

The table VI compares the performance of four optimization techniques: ACO, DE, BO, and PSO, in all the ensemble models used in this paper. From the result, it is clear that PSO achieves the best overall performance with the highest mean accuracy of 0.954, mean F1-Score of 0.979 and best accuracy of 0.956, making it the most effective technique. ACO, while excelling in Mean Recall with a value of 0.978, has the lowest Mean Precision at 0.924 and shows greater variability with a *B*_*A*_-*M*_*A*_ of 0.005 and a standard deviation of 0.005, indicating an imbalance between precision and recall. In contrast, DE and BO exhibit the highest stability, both having the lowest *B*_*A*_-*M*_*A*_ and a standard deviation of 0.001, making them the most consistent techniques. BO slightly outperforms DE in Mean Accuracy, with values of 0.949 compared to 0.946, along with other metrics. PSO is the best choice for maximizing overall performance, while BO or DE are more suitable for scenarios that require stability and consistency.

**TABLE VI.**
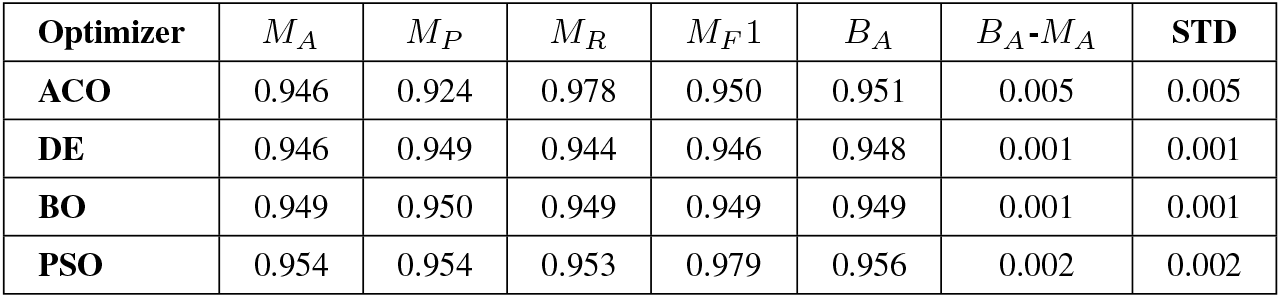
Statistical Analysis of Optimization Techniques across all Ensemble Models.

Figures 16, 17 show that *S*_*E*_ and PSO emerge as the top performers in their respective categories, delivering the best overall precision and the F1 score. However, *SV*_*E*_ and BO or DE are preferable in scenarios where stability and minimal variability are critical. ACO, despite its strong recall performance, shows limitations due to its imbalance between precision and recall.

**Fig. 16.**
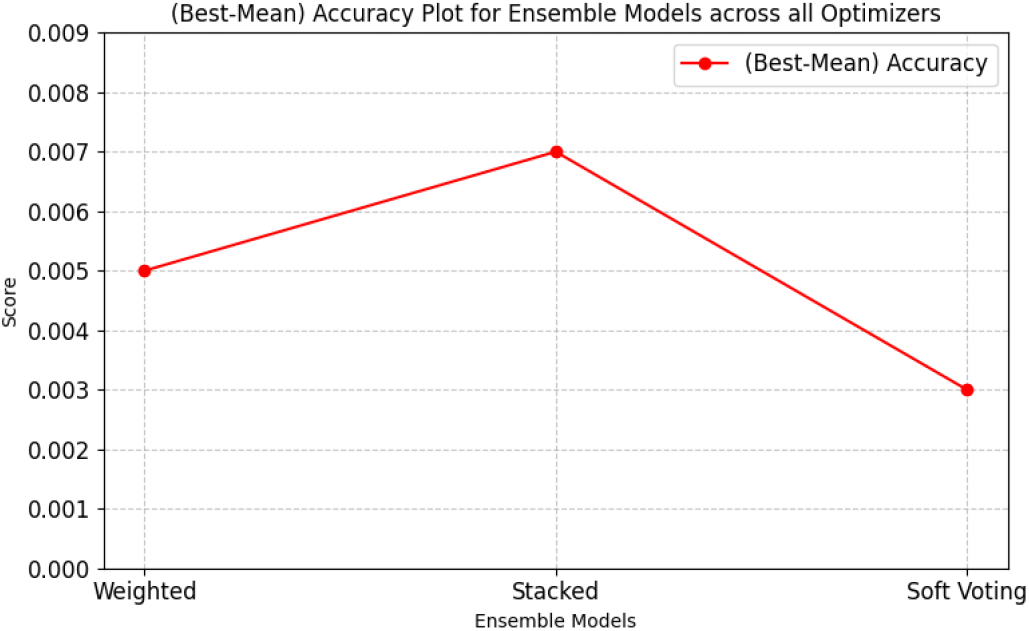
(Best-Mean) Accuracy Comparison for Ensemble Models.

**Fig. 17.**
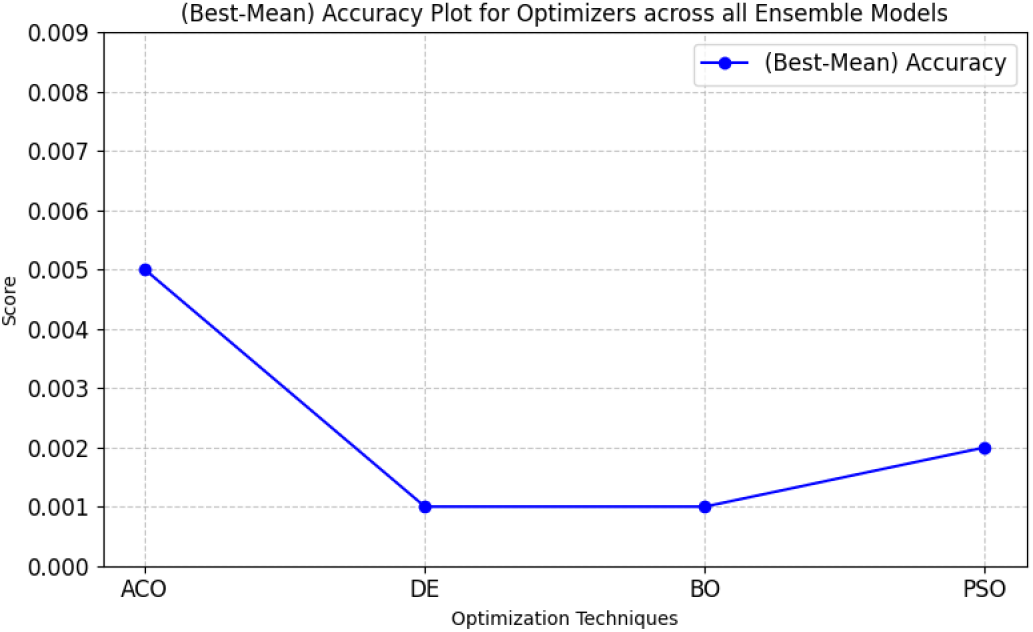
(Best-Mean) Accuracy Comparison for Optimization Techniques.

## VII. Conclusion and Future Scope

The research aims at improving stroke prediction using machine learning techniques, notably through a combination of SMOTEEN data balancing and the PSO optimized stacked ensemble model. Our Proposed Approach demonstrated a high prediction accuracy of 95.6%, highlighting its effectiveness in managing imbalanced datasets. A key focus of the study is to address the bias in machine learning predictions caused by unbalanced training data and disparities in healthcare access. The study argues that SMOTEEN Data Balancing in conjunction with ensemble methods, such as stacking, can help mitigate these biases by improving the resilience and accuracy of the system. We have also compared our proposed approach with Other Ensemble Methods and optimization techniques and found that PSO with Stacked Ensemble is outperforming all other combinations. Future work can explore the integration of deep learning models, such as CNNs or RNNs, to automatically extract complex features and improve prediction accuracy. Combining deep learning with PSO-optimized ensemble methods could further improve performance. Furthermore, deploying and validating these models in real-world settings and developing explainable AI techniques for deep learning would ensure robustness, transparency, and fairness across diverse populations.

## Declaration

### Ethics approval and consent to participate

Not applicable.

### Consent for publication

Not applicable.

## Competing interests

The authors declare no competing interests.

### Author Contributions

A.J., A.K.D., M.G., S.Y., A.P. worked on Writing – original draft, Validation, Project administration, Methodology, Data curation, Conceptualization, while R.A., B.L., S.M. conducted Writing – review & editing, Visualization, Project administration, Data curation.

## Availability of data and materials

The datasets used during the current study are available in Kaggle repository, https://www.kaggle.com/datasets/ fedesoriano/stroke-prediction-dataset.

